# Cortical beta power reflects a neural implementation of decision boundary collapse in speeded decisions

**DOI:** 10.1101/2023.01.13.523918

**Authors:** Hans Kirschner, Adrian G. Fischer, Claudia Danielmeier, Tilmann A. Klein, Markus Ullsperger

## Abstract

A prominent account of decision-making assumes that information is accumulated until a fixed response threshold is crossed. However, many decisions require weighting of information appropriately against time. Collapsing response thresholds are a mathematically optimal solution to this decision problem. However, our understanding of the neurocomputational mechanisms that underly dynamic response thresholds remains very incomplete. To investigate this issue, we used a multistage drift diffusion model (DDM) and also analysed EEG beta power lateralization (BPL). The latter served as a neural proxy for decision signals. We analysed a large dataset (n=863) from a speeded flanker task and data from an independent confirmation sample (n=119). We show that a DDM with collapsing decision thresholds, a process where the decision boundary reduces over time, captured participants’ time-dependent decision policy better than a model with fixed thresholds. Previous research suggests that BPL over motor cortices reflects features of a decision signal and that its peak may serve as a neural proxy for the decision threshold. Our findings offer compelling evidence for the existence of collapsing decision thresholds in decision-making processes.

## Introduction

We often make decisions under time pressure, precluding extensive evidence accumulation in favour of or against choice alternatives. For example, in the realm of financial trading, quick decision-making is paramount. A trader might start the day with a high threshold for certainty, seeking clear signals that a trade will be profitable. However, as the trading day nears its end, the trader may lower this threshold, opting to make trades with less certainty to avoid missing potential market gains. This scenario mirrors the collapsing decision thresholds concept we investigate in our research. Psychology and neuroscience have successfully utilized sequential sampling models such as the drift-diffusion model (DDM) to understand the underpinnings of decision-making (Ratcliff et al., 2016). DDMs assume that motor executions of decisions are triggered when the accumulated evidence for one option/alternative crosses a response threshold. Traditionally, diffusion models assume a time-invariant decision policy, which is reflected in fixed decision boundaries. However, as in the example above, this assumption may be ill-posed on situations where decisions must be made under time pressure. Consequently, more recently, diffusion models with dynamic decision bounds gained popularity (Hawkins, Forstmann, et al., 2015; Palestro et al., 2018; Ratcliff et al., 2016). Here, as time passes, decisions are triggered at decreasing decision thresholds. This facilitates terminating decision making when no solution can be reached within the desired time frame.

Neurophysiological signals that implement time-dependent decision policies have recently been demonstrated in multiple species in the form time-dependent build-up of movement-selective activity consistent with an additive urgency signal. This can be considered as a neural implementation of dynamic decision bounds (Hanks et al., 2014; Kelly et al., 2021; Murphy et al., 2016; Steinemann et al., 2018; Thura & Cisek, 2016), yet our understanding of the underlying mechanisms of such adaptations remains incomplete. It has recently been suggested that the motor cortex is involved in the decision-making process by continually sampling information in favour or against response options (Cisek & Kalaska, 2005; Pape & Siegel, 2016). In the EEG, this process is reflected in beta band (13-25 Hz) activity over centroparietal electrodes. It has been shown that beta power over the contralateral motor-cortex decreases before any overt action and even reflects imaginary movements (Kuhn et al., 2006). The difference in beta power between both motor-cortices may thus encode the relative evidence in favour of respective response options in a lateralised fashion (Donner et al., 2009; Pape & Siegel, 2016), or, put more broadly, decisions in action-space (Cisek & Kalaska, 2005; Hunt et al., 2013). In this line of research, we recently demonstrated that model parameters of a multistage drift-diffusion model and the lateralisation of EEG beta power convergently show a complex interplay that facilitates behaviour adaptations after erroneous responses in a flanker task (Fischer et al., 2018). Specifically, suppression of distracting evidence, together with increased response thresholds appear to cause slower and more accurate post-error responses. Based on this work demonstrating a remarkable similarity of the time course of the decision variable in the DDM and the beta power lateralization (BPL) over motor cortex, we argue that the BPL peak around response time can serve as a proxy for response thresholds.

Here, we follow up on this argumentation and investigate whether BPL can demonstrate a plausible neural mechanism for collapsing decision thresholds in a large sample of 863 healthy participants. We complemented our previous multistage DDM (Fischer et al., 2018) by an additional free parameter that allowed decision bounds to collapse according to a cumulative Weibull distribution. As depicted in Figure 1D, this model makes distinct predictions about the decision threshold depending on speed: later decisions are made at a lower threshold. We hypothesized that if decision thresholds collapse, a decreased BPL peak for slower responses should be present.

**Figure 1.**
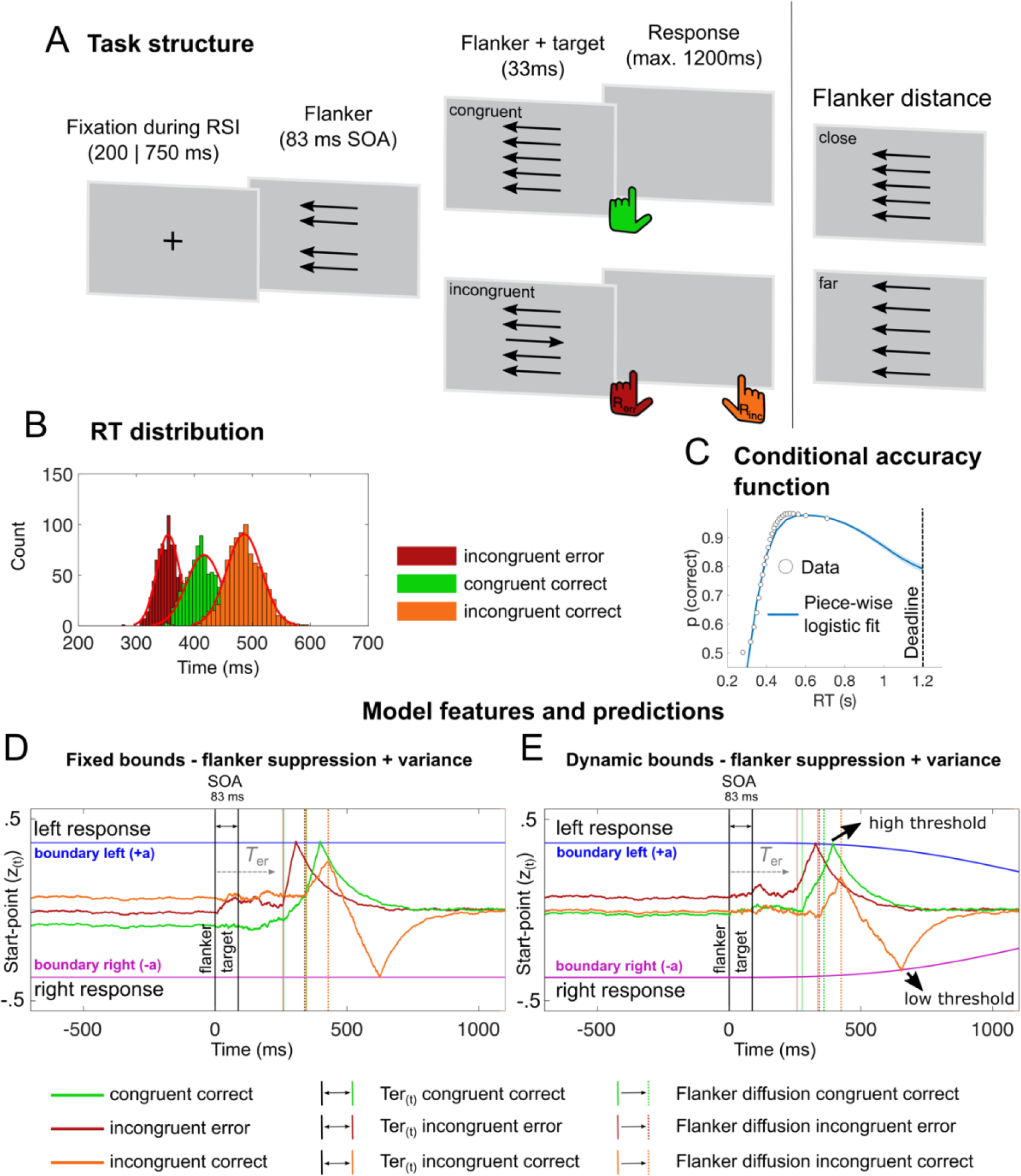
Task information, model features and predictions. **(A)** In the flanker task either congruent or incongruent flankers (four surrounding arrows) in proximity or further away from the central target (inlet) are presented on each trial. Congruent stimuli usually yield faster responses (green) compared to incongruent stimuli, which lead to delayed responses (orange) or induce fast errors (red) **(B)**. **(C)** Shows the conditional accuracy function. Here, points indicate mean accuracy of trials sorted by RT into 25 equal-sized bins. The line shows best fits of piecewise logistic regressions to each subject’s single-trial data. Shades indicate ± s.e.m. **(D-E)** Illustration of the multi-stage DDMs based on simulated single trial decision processes. On a given trial, the decision can be randomly biased to favour one response (startpoint variance, sz_(t)_). The first stage of the respective model consisted of a pre-stimulus zero-mean baseline for which the value is determined by the start-point (z_(t)_) (Stage 1). After flanker presentation (0 ms), visual processing and motor execution times are captured by the non-decision time (T_er_) parameter, which can vary from trial to trial (st). In this period the decision process randomly drifts away from the start point (Stage 2). The next stage represents a noisy diffusion in flanker direction (f_(t)_) for the duration of the SOA which again can vary from trial to trial (sf) (Stage 3 - flanker diffusion). After flanker diffusion, stage 4 models the consecutive diffusion into the correct response direction with drift rate v_(t)_ with trial-to-trials variance (sv). Longer RTs for incongruent correct trials are due to an initial drift away from the correct response (orange decision process). Error likelihood increases when the baseline is shifted towards the flanker direction and f_(t)_ is higher on a given trial resulting in an early crossing of decision boundaries (red decision process). As a result, errors are usually faster on incongruent trials. Finally, Stage 5 models the return to baseline of the decision variable, akin to an Ornstein-Uhlenbeck process. The height of the boundary parameter (a) determines how much evidence accumulation is required to cross the boundary and trigger a response. Which can either be fixed **(C)** or collapse with increasing decision time **(D)**. To simulate the temporal evolution of the single trial decision process depicted in **(C)** and **(D)** we used the mean maximum likelihood parameters from the group fit obtained for DDM3 and DDM4.

We found that the extended DDM provides a better fit for the data. This model predicts decision threshold differences due to boundary collapse over time. This is confirmed in BPL, where slower responses show a decreased peak. These data suggest that peak BPL around response time may serve as a neural proxy for the decision threshold. Taken together, model predictions and their independently measured neuronal proxies in beta power convergently support the assumption of collapsing decision thresholds.

## Results and Discussion

863 healthy subjects performed a speeded arrow-version of an Eriksen flanker task while EEG was recorded. Participants were instructed to respond as quickly and accurately as possible to the direction of a centrally presented arrow (see Figure 1A). To elicit a large proportion of response errors, we presented distracting flanking stimuli in visual proximity and slightly ahead in time (83 ms) of the imperative, central target stimulus. On incongruent stimulus sets a mismatch between the flanker and target arrow directions induced conflicting response tendencies. The response conflict was further enhanced by manipulation of stimulus distances among each other (close and far, Danielmeier et al. (2009)) and response-stimulus intervals (short and long, Danielmeier and Ullsperger (2011)). To increase time-pressure, we kept the response stimulus intervals (RSIs) and the response window of 1200 ms rather short. Behavioural analyses of this task confirmed that general behaviour effects were in accordance with typical results seen in flanker tasks. They reflected interference effects, with slower (ΔRT = + 62 ms, t(862) = −81.79, p < .001, d = 1.03, BF_10_ > 100) and more erroneous (Δaccuracy = 20%, t(862) = 96.18, p < .001, d = 3.44, BF_10_ > 100) responses on incongruent trials, whereby these effects interacted with flanker distance. Moreover, all error-related behavioural adaptations associated with cognitive control (post-error slowing (PES), post-error increase in accuracy (PIA), post-error reduction of interference (PERI)) were present. Overall participants responded accurately (mean accuracy > 85%) and most errors were committed at faster RTs compared to correct responses (see Fischer et al., 2018 for a detailed description of these results). Importantly, we investigated time-dependency in the decision process utilizing empirical conditional accuracy functions (CAF) similar to Murphy et al. (2016). CAFs relate accuracy to RT. A time-dependent decision policy has been characterised by a combination of a few missed deadlines, negative CAF slopes and decreased accuracy around the response deadline (Murphy et al., 2016). We employed single-trial logistic regression to estimate the shape of the empirical CAFs (see Figure 1C and methods). After accounting for fast response errors, the estimated CAF slope was negative for slower responses. Moreover, using the regression fit to estimate the accuracy around the deadline revealed decreased accuracies. Finally, the amount of missed deadline was low: 3.49 % (SD: 3.59). Combined, these findings indicate that participants employed a time-dependent decision policy.

### An extended DDM with collapsing decision boundaries captures task behaviour best

We have previously shown that an extended multistage conflict DDM (depicted in Figure 1D and E) captures these behavioural effects (Fischer et al., 2018). This model assumes that choice options are mutually exclusive. Therefore, a single decision variable is reflecting both decision options. On every trial, the decision can be randomly biased slightly to one response option (startpoint variance, sz). The height of the boundary parameter (a) determines how much evidence accumulation is required to cross the boundary and trigger a response. The non-decision-time-parameter (*T*_er_), reflects the time from actual flanker onset until the information can be used to influence the decision process, which in our model can vary from trial to trial (st). During flanker-only presentation, evidence was accumulated in the direction of the flankers (83 ms). After target onset, evidence accumulation is driven by the direction of the target. We previously demonstrated that a model that allowed for trial-by-trial weighting of the distractor-driven diffusion (simulating an attentional process suppressing the influence of the flankers) explained participants’ behaviour best (Fischer et al., 2018). Comparisons of model estimates fitted to either post-error or post-correct trials revealed that post-error adaptations are facilitated by a complex interplay of multiple mechanisms: suppression of distracting evidence, response threshold increase, and reduction in the speed of evidence accumulation.

Conceptually, our DDM is very similar to previously introduced DDMs that model behavioural effects in conflict task suggesting that drift rates vary over time (e.g., the dual-stage (Hubner et al., 2010) or the shrinking spotlight model (White et al., 2011)). Here, we use a simplified version of these DDMs, that only assumes that attention shifts from flankers to the target once the target is on screen and that these two stages of information processing can differ from each other (f) which should be different on each trial (sf). Therefore, we did not have to simulate two processes (stimulus and response selection - dual-stage (Hubner et al., 2010))) nor assume a shrinking function for the attentional spotlight (shrinking spotlight model (White et al., 2011)). In addition, our task design differs slightly from the designs used by Hubner et al. (2010) and White et al. (2011) such that flanker stimuli alone precede the presentation of the full target-flanker stimulus array which renders our simplification plausible. Moreover, it should be noted that we do not exclude that separate racing accumulators are a valid model of speeded decision-making under conflict. Yet, we previously demonstrated that the approximation in our DDM which assumes that evidence for response A counteracts evidence for response B, is valid and compatible with the neural signal reflected in BPL (Fischer et al., 2018).

To investigate whether dynamic decision boundaries explain subjects’ behaviour better, we complemented our multistage DDM by an additional free parameter that scaled decision bound collapse according to a cumulative Weibull distribution (see Supplementary Figure 1 for illustration of different boundary dynamics). As can be seen in Figure 1D, this model makes a specific prediction: later decisions are formed at a lower threshold. Model comparison (Figure 2C - L) revealed that our data is better described by the collapsing bound model than by the original models with fixed bounds (as measured by lowest approximated BIC and achieving protected exceedance probabilities of 100%). This indicates that individuals apply a discounting function that decreases decision thresholds when time is running out (i.e., the response window of 1200 ms closes). Sufficiency of the winning model was evaluated through posterior predictive checks that matched behavioural data on various summary measures. Specifically, the model captured correct vs. error RT quantiles for congruent and incongruent trials and conflict related accuracies (Figure2 G-J). Additional model validation analyses indicated that the collapsing bound model tended to recover parameters that generated synthetic data (see Supplementary Figure 3).

**Figure 2.**
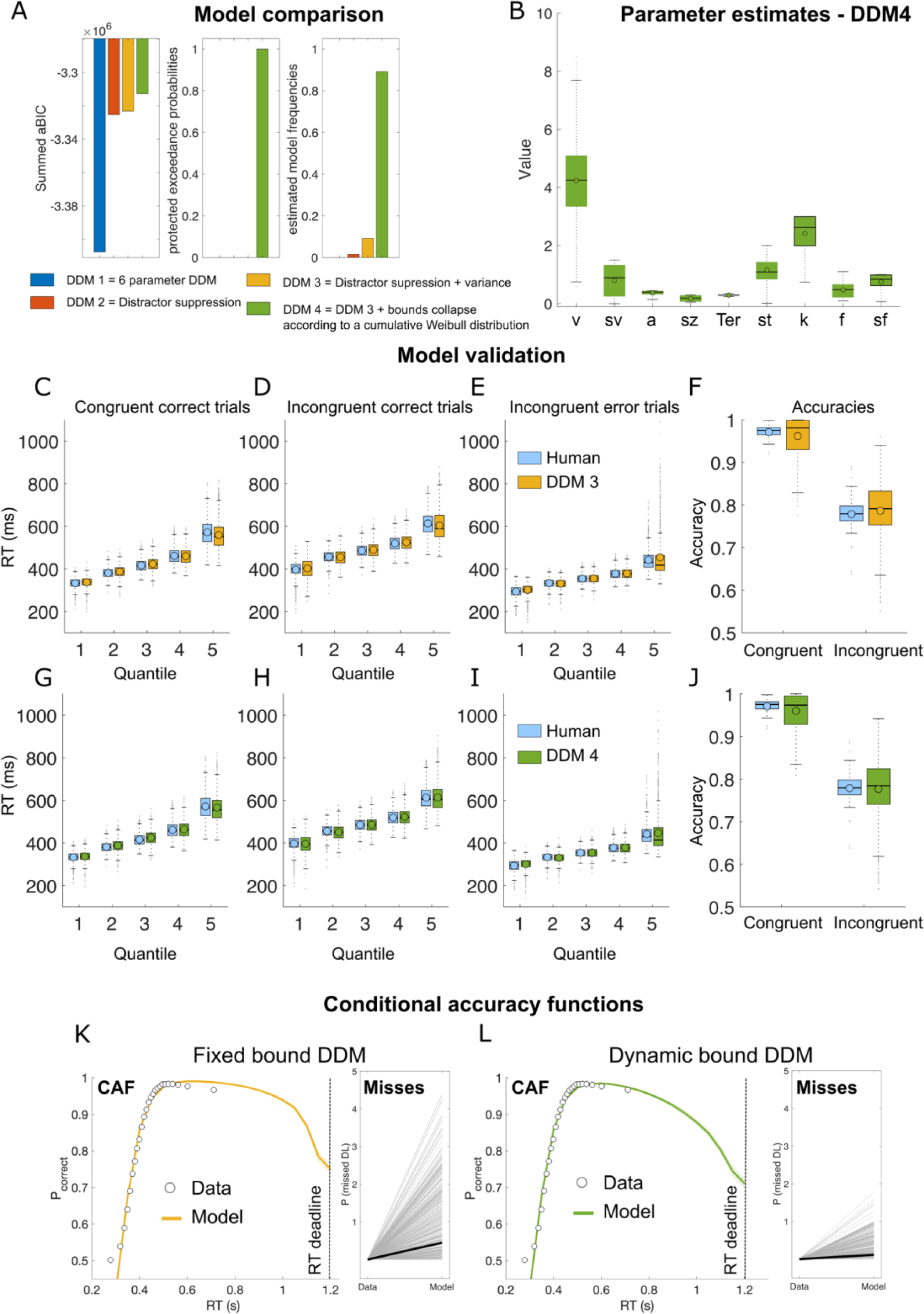
Drift-diffusion model fits and model validation. **(A)** Cumulated approximated Bayesian Information Criterion (aBIC) scores over participants of each candidate model. Higher values indicate a better fit of the models to the behavioral data. Results indicate that the dynamic bound model (DDM4) provides the best fit to the data. Protected exceedance probabilities (pEP) similarly favor DDM4. **(B)** The collapsing boundary model included nine free parameters. In addition to the standard-parameters (DDM3) reflecting drift rate (v), flanker weighting (f), boundary (a), and non-decision time (Ter), and variance parameters that determined how much standard-parameters varied from trial-to-trial (sv, sz, st, sf), the dynamic boundary model included the free parameter k that scaled the boundary collapse that increases with the duration of the decision See Supplementary Table 1 for more details on the Maximum likelihood parameters of three best fitting DDMs. **(C – J)** shows quantile fits of the model (**C-E** DDM3, orange; **G-I** DDM4, green) against human RT data (light blue) and **(F and J)** shows model and human accuracy (**F** DDM4; **I** DDM3). In all conditions (congruent & incongruent correct as well as incongruent error), both models capture the RT data in each quantile, suggesting a good fit to the data. However, the collapsing boundary model appears to capture the variance in the data better (particularly in the later RT quantiles). **(B - J)** boxes = interquartile range (IQR), o = median, - = mean, whiskers =1.5 × IQR, grey dots = outlier. Differences in model fits become more evident comparing CAFs and missed deadlines produced by the model. **(K-L)** depicts Conditional accuracy functions and proportion of missed deadlines from the full fixed bound model (DDM3; **K**) and the collapsing boundary model (DDM4; **L**). Points indicate mean accuracy of trials sorted by RT into 25 equal-sized bins informed form the empirical data. Line shows best fits of piece-wise logistic regressions to each subject’s modelled single-trial data. Shades indicate ± s.e.m. Note that for model fitting we excluded all missed trials from the empirical data.

While the full fixed boundary model (DDM3) also provided a good fit to the data (Figure2 C-F), the collapsing boundary model appears to capture the human data slightly better in later quantiles. This difference becomes more evident in the CAFs that were fit to the modelled data (Figure2 K-L). Here, the collapsing bound model captures the empirical data better. The poorer fit of the fixed bound model may be due to the fact that it is unsuited to generate negative CAF slopes without increasing the number of missed deadlines (Murphy et al., 2016). To lower the CAF slope the fixed bound model must increase the drift-rate variance, which increases the number of missed deadlines.

To characterise predictions of the collapsing boundary DDM we simulated 5000 individual trials using the mean of the best fitting individual parameters (see Supplementary Table 1). When averaged across trials in a median-split analysis, the decision signal shows clear response threshold differences, whereby fast responses show a higher threshold (Figure 3E). Interestingly, our results are at odds with previous research identifying evidence in favour of standard DDMs with fixed bounds in humans (Hawkins, Forstmann, et al., 2015; Hawkins, Wagenmakers, et al., 2015; Ratcliff et al., 2016). In these studies, other decision-making tasks were used (i.e., random dot motion, brightness discrimination, dot separation) and DDMs treat (perceptual) interference trials as trials on which drift rates are low to capture the increased RT and uncertainty on (error)-trials. However, lower drift rates per trial cannot produce fast errors. Yet, fast errors are a hallmark of speeded reaction time tasks, like our flanker task. Here, fast errors arise from an interaction of distractor-driven biasing and trial-wise pre-lateralization of the decision variables as well as noise during stimulus processing (Fischer et al., 2018). Moreover, in contrast to most other decision-making tasks, after flanker and target presentation (which was only presented for 33 ms in our task) no further information could be sampled to inform decisions. Thus, decision policies might be task-dependent and affected by various factors. It is also possible, that mild collapsing decision boundaries were present in previous investigations, but not clearly identifiable in model fit comparison (Murphy et al., 2016). Specifically, Murphy et al. (2016) argue that several popular fixed thresholds DDMs include variability parameters, that are able to capture similar behavioural effects as in dynamic boundary DDMs (e.g., Brown & Heathcote, 2008; Murphy et al., 2014), which precludes discrimination between the two model classes. Indeed, our fixed boundary model (DDM3) included variance parameters and captured subjects’ behaviour well. On average, the boundary collapse in DDM4 was mild (see Figure 1E) and, hence, only affected a small percentage of trials. The differences between fixed and dynamic boundary models therefore may exert a hard-to-detect influence on likelihood estimates in small samples and with methods commonly used for model comparison (e.g., summed BICs). Yet, predictions of different sequential sampling model classes can be highly informative to differentiate between different mechanistic accounts. In this line of thought, we also considered a different kind of dynamic decision model. In this model, decision making under time pressure is facilitated by an urgency signal (Cisek et al., 2009; Hanks et al., 2014). Here, an urgency signal that increases with decision time is used as a multiplicative gain of the diffusion and noise (DDM5; see Supplementary Figure 1 and 2A for an illustration). Both dynamic DDMs capture RT quantiles and accuracies better than the time-invariant models (Supplementary Figure 2B) but make distinct predictions regarding the response threshold. While the collapsing bound model predicts that slower responses are made at lower decision boundary thresholds, the urgency model would predict similar decision boundaries for slow vs. fast responses (Supplementary Figure 2D). We test the neural plausibility of these accounts in the next section.

**Figure 3.**
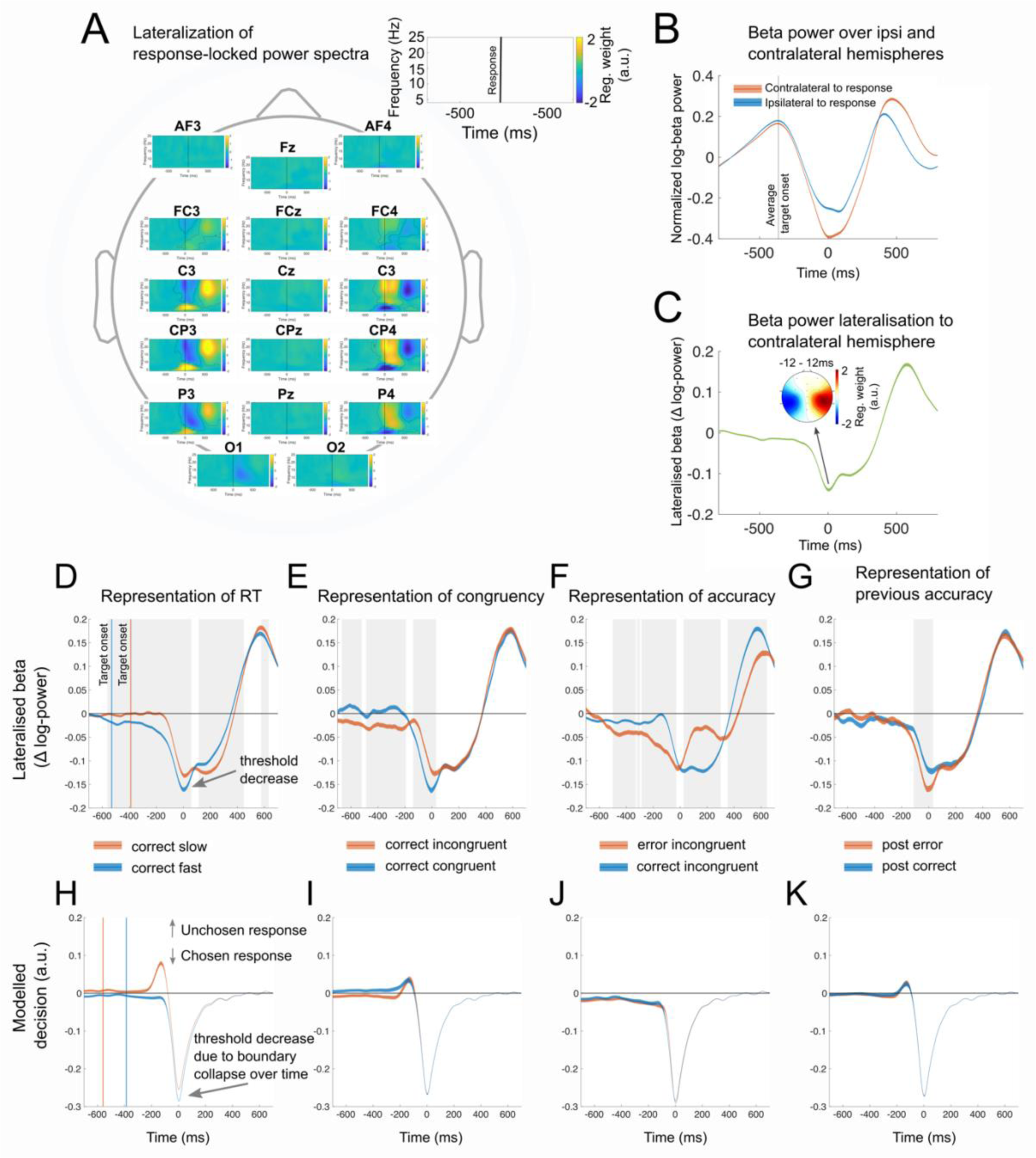
Response-locked power spectra, beta power, and DDM decision variable. **(A)** Shows the result of a single-trial regression analysis comparing responses with the left and right hand, while controlling for congruency, flanker distance, response stimuli intervals (RSI), following RSI, trial number, and log transformed RT. Data suggests, that Beta power more strongly decreases when a response is given with the contralateral compared to the ipsilateral hand. This effect was most pronounced over the electrodes C3/4 and CP3/4. Beta power lateralization signal **(C)** is composed of the activity-difference recorded over contralateral and ipsilateral motor cortices **(B)**. **(D - G)** shows neural stimulus processing locked to response separated by response speed, congruence, accuracy, and previous accuracy. First, we show that in BPL as well as modelled signal a lateralisation towards the given response is pronounced for fast response **(D & H)**. Importantly, BPL at response execution is reduced for later responses. This observation confirms dynamically changing decision bounds within a trial and is also reflected in the modelled signal **(H)**. In addition, when comparing RT-matched correct congruent and incongruent trials, we found a significant difference at response execution **(E)**. Exploratory analysis of the lateralised signal (see Supplementary Figure 4) demonstrated that this effect is caused by reduced activation (i.e., increased beta power) of the ipsilateral hemisphere on congruent trials compared to incongruent ones. This likely reflects enhanced motoric readiness caused by the flanker stimuli. This effect is not captured by the DDM signal **(I)** because it does not include threshold differences between trials. When comparing RT-matched correct and error responses **(F & K)**, we demonstrate that both conditions are associated with similar thresholds in BPL. This effect is similarly reflected in the modelled DDM signal. Finally, we demonstrate that BPL at response execution is increased on response matched post error trials **(G)**. This may reflect strategic decision policy adaptations to increase accuracy after errors (Fischer et al., 2018; Murphy et al., 2016). Again, this effect is not reflected in the modelled DDM signal because this model does not include threshold differences between trials **(K)**. *Note.* Contours in **(A)** represent significant clusters after cluster-based permutation testing (Maris & Oostenveld, 2007). Shades in **B-K** represent 99.9% CIs. Grey backgrounds in indicate significant time-points after Bonferroni correction. In **H-F** we rectified the modeled diffusion signal to always plot the unchosen response up and the chosen response down. For details about the simulation see Methods.

### Cortical beta power lateralization reflects neural implementation of decision boundary collapse

Next, we tested the hypotheses that the response-locked BPL represents a neural implementation for decision boundary collapse. First, we confirmed that beta power decreased more at centro-parietal electrodes contralateral to the initiated response (see Figure 3A-C), suggesting that BPL reflects differential motor activation (Donner et al., 2009; Pfurtscheller, 1981). When averaged across trials in a median-split analysis, differences in BPL are clearly evident in the response locked data, where fast responses show a higher BPL (i.e., higher decision boundary; fast RT: −0.13 (SD: .003) vs. slow RT: −0.10 (SD: .003); t(862) = −9.58, p < .001, d = 0.25, BF_10_ > 100, see Figure 3D). The representation of RT is similarly reflected in the modelled DDM signal (Figure 2H). Thus, collapsing boundaries are confirmed by response-locked BPL, indicating that humans make use of a discounting function to decide earlier to fulfil a time criterion. To confirm this finding on a single-trial level we regressed task-related behavioural factors onto measured individual single-trial BPL at response execution (button press ±12ms). This allows to control for other task factors that may additionally influence BPL but are not of primary interest for this report (but see Supplementary Figure 4 for more details on these results). The regression analyses confirmed that BPL around the response decreased (i.e., boundary collapse) with increasing RTs (t(862) = 13.26, p < .001, d = 0.45, CI [0.46, 0.62], BF_10_ > 100, see Figure 4A). Plotting the RT quantiles against the BPL at response execution confirms a boundary collapse that closely resembles a sigmoidal function (Figure 4B). Yet, a correlation between BPL and RT by itself does not necessarily imply bound collapse on our task. Specifically, trials with slower RTs are likely to be ones in which there was greater response conflict and so one could predict greater build-up for the ultimately unchosen alternative and an overall reduction in lateralisation. To address this issue, we conducted several control analyses. First, our GLM consisted of various task factors including congruency to control for confounds and the interdependence of effects. Second, in an explorative analysis, we split the regression analysis by ipsilateral (hemisphere that does not induce the choice on the current trial) and contralateral hemispheres to better understand which hemisphere drives RT-related BPL decreases (see Figure 4C). Results indicate that threshold decreases are partly due to slightly stronger decreases in beta power over the contralateral hemisphere (contralateral: t(862) = 5.13, p < .001, d = 0.17, CI [.19, .42], BF_10_ = 1.57e^+4^ vs. ipsilateral: t(862) = −3.68, p < .001, d = 0.12, CI [−.33, −.10], BF_10_ = 32.10). However, the effect is stronger when the relative signal per trial is investigated, than when either of the hemispheres beta power is analyzed alone (contrast contralateral against BPL effect: t(862) = 3.87, p < .001, d = 0.23, BF_10_ = 63.47). These data suggest that peak BPL, coinciding with the motor response, may be an especially valid marker of response threshold modulations. However, it should be noted, that this assumption may be limited to speeded forced-choice tasks with short reaction times and evidence accumulation (Rogge et al., 2022). In the averaged beta power signal, this effect is partly explained by a reduction of beta power over the contralateral hemisphere (Figure 4D), while on average beta power decrease over the ipsilateral hemisphere is barely changed when responses are given later in a trial (Figure 4E). A closer look at beta power around the response (button press ±12ms) divided into five equally sized RT bins indicates that both beta power over the contralateral and ipsilateral hemisphere to the response hand are RT dependent (Figure4 D&E). Here, RT dependency in beta power amplitudes follow an inverted U-shaped curve. Considering previous research that characterised peak beta-power contralateral to the response as “motor threshold” (e.g., Kelly et al., 2021), these data suggest that fast response errors partly arise from decreased motor thresholds (see Fischer et al., 2018 for a detailed discussion on how fast response errors in simple tasks arise) and that collapsing decision boundaries terminate decision-formation under time pressure, which potentially increases error likelihood. This pattern of results implies that beta power around response execution is fairly variable, which adds to the accumulative evidence of showing time-dependent effects on beta amplitudes (see discussion below). Critically, our results suggest, that peak BPL (i.e., the relative beta signal), coinciding with the motor response, may be an especially valid marker of collapsing decision thresholds, as it incorporates both, contra- and ipsilateral RT dependent motor preparation signals.

**Figure 4.**
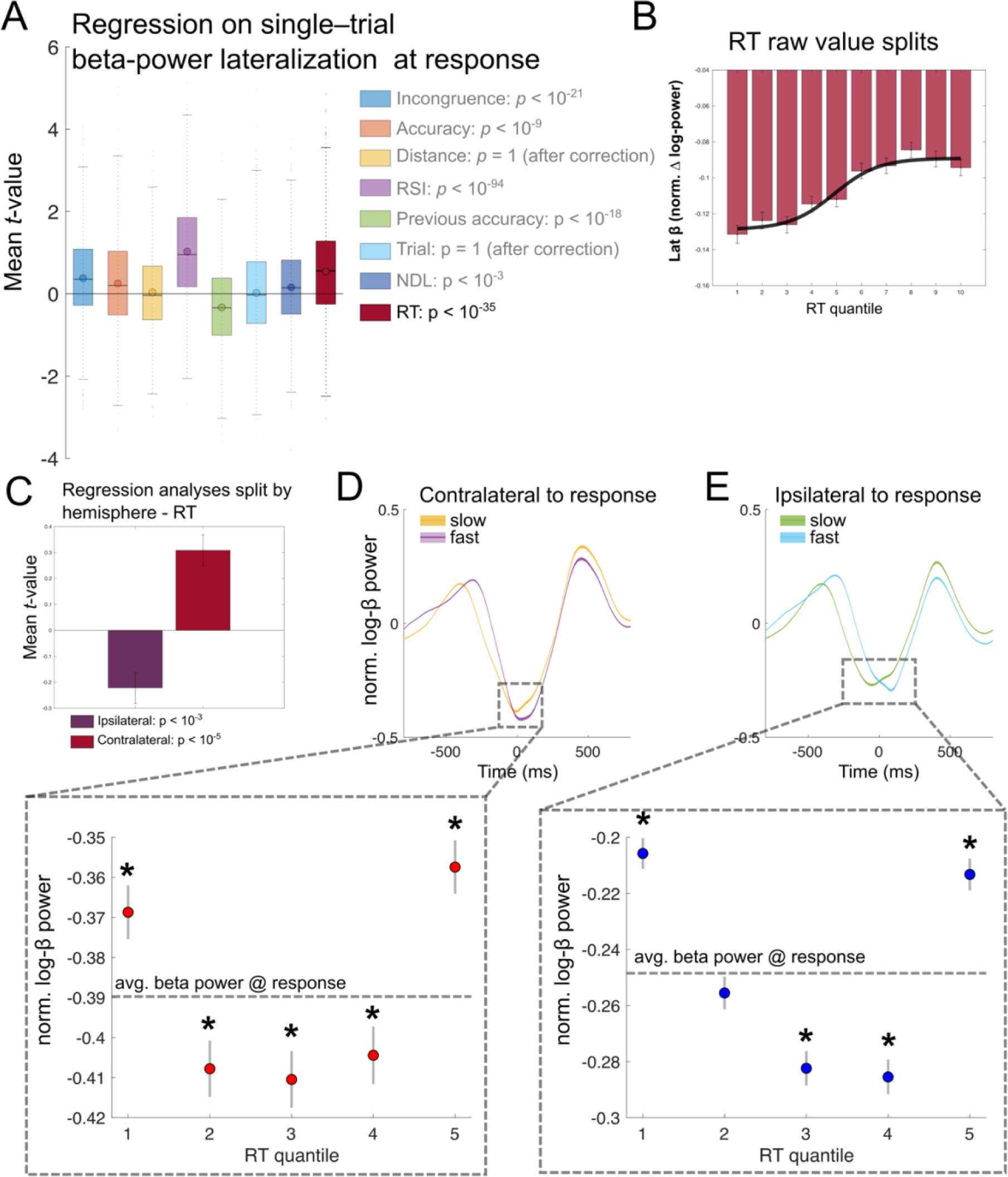
Single-trial regression on beta-power lateralization at response. **(A)** Results of a single-trial regression analysis on BPL at response (mean ± 12ms surrounding button press). This analysis confirms collapsing decision boundaries on a single-trial level while controlling for other task factors that potentially influence BPL. **(B)** BPL raw values for RT quantiles plotted against BPL at response. Fitting the mean values trajectory confirms boundary collapse that closely resembles a sigmoidal function (black line expressed with: ymin = −0.13, ymax = −0.09, x0 = 4.89, and slope = 0.49, adjusted R^2^ = .95). **(C)** Display results of separate regression analyses on beta power split by ipsilateral (hemisphere that does not induce the choice on the current trial) and contralateral hemispheres. RT related BPL decrease is partly due to slightly more decreases over the contralateral hemisphere. In the averaged beta power signal, this effect is partly explained by a reduction of beta power over the contralateral hemisphere **(4D)**, while on average beta power over the ipsilateral hemisphere is barely changed when responses are given later in a trial **(E)**. Boxes below show beta power (mean ± 12ms surrounding button press) divided into five equally sized (within subject) RT bins. * = uncorrected significant difference to mean beta power (dashed lines) **(A)** boxes= interquartile range (IQR), o = median, - = mean, whiskers =1.5 × IQR, grey dots = outlier. Error bars reflect standard error of the mean. **(A and C)** depicts mean within participants t-values, and statistics are results of t-tests of individual within-subject regressions against zero. RSI reflects the response-stimulus interval, NLD the distance from the last break in the task, and trial the general time on task.

### Specificity of the effects to the beta frequency range

Similarly to BPL, lateralization in the alpha (8 −12 Hz, APL) and μ (8–14 Hz, MuPL) frequency band over the motor cortex has been related to a variety of decision processes, such as decision formation and preparation (e.g., Donner et al., 2009; Pape & Siegel, 2016; Rogge et al., 2022) or urgency of a decision (Murphy et al., 2016). In a control analysis we therefore aimed to establish specificity of our effects to the beta frequency range. First, we confirm that sensorimotor alpha and μ power similarly reflected response-related signals and that this was specifically lateralised to the active motor cortex (see Figure 3A and Supplementary Figure 5). Yet, effects are smaller and peak slightly after responses are made (Supplementary Figure 5B and C). To quantify the effect differences on a single-trial level, we regressed response speed onto measured individual single-trial APL and MuPL at response execution while controlling for other task factors (see Supplementary Figure 6 for details). Results revealed that, response speed is similarly reflected in movement-selective alpha and *μ* suppression, but that this effect is more pronounced in the beta frequency range (BPL vs. APL: .54 (.04) vs. .23 (0.04); t(862) = 6.43, p < .001, d = 0.26, BF_10_ > 100; BPL vs. MuPL: .54 (.04) vs. .28 (.04); t(862) = 6.15< .001, d = 0.26, BF_10_ > 100). In addition to the alpha and μ frequency band, an opposite effect was present for lateralization in the theta frequency band (4–8 Hz) with a similar topography (see see Figure 3A and Supplementary Figure 5 and 6), which is not further interpreted in this report.

### Conclusion and implications for beta power function in the decision process

This study uncovers compelling evidence to suggest that under high time pressure, dynamic decision thresholds drive the termination of decision-formation, as seen in the beta power lateralization (BPL) over the motor cortex during a speeded Flanker task. This is demonstrated by behavioral modelling, showing that subjects time-dependent decision policy is best captured in DDMs that allow dynamic decision bounds and confirmed in the neural signal, whereby BPL over motor cortices reflects features of the modelled decision signal.

Taken together, our results extend previous work on BPL as a neural signal that carries important features of a decision variable (Donner et al., 2009; Fischer et al., 2018; O’Connell & Kelly, 2021). Recent work has suggested that beta power suppression around response contralateral to the chosen hand reaches a highly similar level, irrespective of urgency or response speed, suggesting a fixed response threshold (Corbett et al., 2023; Feuerriegel et al., 2021; Kelly et al., 2021; Murphy et al., 2016; Steinemann et al., 2018). In line with this account, it has been suggested the beta power over the right and left motor cortex is reflecting two race-to-(response-)threshold motor preparation signals (Corbett et al., 2023; Kelly et al., 2021; O’Connell et al., 2012). Under time-pressure, speeding may be facilitated by an urgency signal that independent but in addition to evidence, drives the signal towards the response threshold (Cisek et al., 2009; Corbett et al., 2023; Hanks et al., 2014; Murphy et al., 2016; Steinemann et al., 2018). Recent work in non-hum primates (Hanks et al., 2014; Thura & Cisek, 2016) and humans (Murphy et al., 2016) demonstrated that under time pressure the neural urgency signal is characterized as a time-dependent increase in common activation for both the chosen and unchosen response alternatives, which translates into a diminishing contra/ipsi μ lateralization with increasing RT. Our results add to this line of research in several ways. First, to the best of our knowledge, this is the first study to demonstrate a neural implementation of collapsing decision thresholds in a speeded Flanker task. Second, we show that motor response thresholds contralateral to the response are more dynamic than previously thought. Third, we suggest that peak BPL (i.e., the relative beta signal), coinciding with the motor response, may be an especially valid marker of response threshold modulations as it incorporates both, contra- and ipsilateral motor preparation signals.

Interestingly, BPL in our task resembles the characteristic of a signal that dynamically reduces the threshold it must reach to trigger a response. Such a signal has previously been demonstrated in the form a centro-parietal positivity (CPP) in the EEG, which reduces in amplitude alongside RT, accuracy and urgency (Kelly et al., 2021; Steinemann et al., 2018). In an independent confirmation sample (n = 119) collected at a different site, but using the same study protocol, we replicated all key findings (see supplementary results). Thus, in this study we provide strong, converging evidence that movement-selective beta power lateralization reflects a signal that dynamically adjusts response thresholds to terminate decision-formation under time pressure. Our finding that BPL reflects collapsing decision boundaries under time pressure is consistent with the additive urgency signal account. In fact, in an abstract mathematical model both accounts are very related. This was also reflected in a similar fit of the collapsing bound (DDM4) and urgency model (DDM5) to the empirical data in our task. Yet, our finding of time-dependent motor thresholds contralateral to response is surprising. How can this discrepancy in findings be explained? One factor that likely contributes to the differences is the nature of our task. In our task, after flanker (83ms) and target presentation (which was only presented for 33 ms in our task) no further information could be sampled to inform decisions. In contrast, other studies typical use paradigms in which evidence can be accumulated until the response is made (e.g., random dot motion paradigms). Hence, the differences in findings may highlight that decision policies may not be generalized across tasks. Indeed, there is evidence that response policies could be task specific even in variants of conflict tasks like the Simon compared to the Flanker task (Hübner & Töbel, 2019). It is also possible, that within-trial adjustments of response thresholds are small and are more pronounced in relatively rare cases. For example, in our sample very slow errors and slow responses in trials with no visual conflict are sparse. Here, we draw inference from a large sample and likely have the power to detect even small time-depended changes and rare response constellations. In addition, beta amplitudes vary quite widely across groups and thus small effects might be difficult to detect in smaller samples. Indeed, even our confirmation sample is considerably larger than in average EEG studies. Finally, our results do not exclude the possibility of an urgency signal as suggested by Thura and Cisek (2017). It may be, that instead of increasing the gain in the decisions process, the urgency signal in the globus pallidus modulates decision thresholds in the cortex. Yet, this argumentation remains speculative.

In summary, our findings build on previous research on movement-selective beta power. They suggest that one mechanism for achieving fast-paced decisions under time pressure is through collapsing response thresholds, which facilitate dynamic decision policy adjustments according to task demands.

## Materials and methods

### Participants

Eight hundred ninety-five healthy young adults were recruited into the study. After exclusion of participants due to low task performance, recording failures, or poor data quality, 863 participants remained for subsequent analyses. The mean age of the sample including 434 female and 429 male participants was 24.2 years (range: 18–40). A detailed description of the sample characteristics and exclusion criteria can be found in Fischer et al. (2018).

### Task

We employed a speeded arrow version of the Eriksen flanker paradigm in that participants were encouraged to respond as quickly and correctly as possible to the direction of the target arrow presented in the centre of the screen. The target could point either to the right or the left side, requiring a corresponding response with the right or left thumb. We additionally modulated response-stimulus intervals (short = 250ms, long = 700ms) as well as the distance between the imperative central and distracting flanker arrows (far distance: 6.5° and 4°, and close 3.5° and 1.75° visual angle). In total the task comprised 1,080 trials (see Fischer et al 2018, for more details on the task). A short response window of 1200ms and short RSIs ensured high-time pressure.

### Empirical conditional accuracy functions

Analyses of the conditional accuracy functions were conducted using slightly adapted code provided by Murphy et al. (2016). The procedure was as described by Murphy et al. (2016) and the following description is adapted from therein.

To estimate mean accuracy as a function of RT (i.e., the CAF) we used single trial logistic regression. To account for fast errors and for a possible decreasing CAF towards the response deadline, an algorithm was constructed that minimizes the combined sum of squared errors of piece-wise logistic regressions of accuracy (1 = correct, 0 = error) onto RT, splitting trials before and after a temporal inflection point α such that:

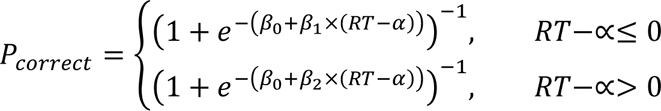

Here, *β*_0_is accuracy at *α*, *β*_1_ is the slope of the CAF before *α*, and *β*_2_ is the slope of the CAF after *α*. *β*_1_ was constrained to be ≥ 0 to reflect the fact that the left segment of the piece-wise fit accounts for the increasing portion of the CAF (i.e., this considers the fact that in our task most response errors are made at fast RTs). The model was fit using Nelder-Mead simplex minimization to estimate *β*_0_, *β*_1_and *β*_2_ parameters while conduction an exhaustive search of possible *α* values (step-size = 10 ms ending at 1s). Whichever piece-wise segment is fit first determines *β*_0_and thus constrains the fit of the remaining segment; therefore, the algorithm was run twice (left segment fit first and right segment fi first) for each *α* to find the true minimum (Karşılar et al., 2014; Murphy et al., 2016). This single-trial regression approach allows for a reasonable approximation of the temporal evolution of the CAF throughout the whole trial length.

### Drift-diffusion model and model comparison

Features of the multi-stage sequential sampling models one to three, and general fitting procedures were exactly as described by Fischer et al. (2018). The description is adapted from therein. We used a multi-stage sequential sampling model to simulate participants’ RT and accuracy distributions and the decision process giving rise to these. This discrete diffusion model simulates decisions as a Wiener process with stepwise increments according to a Gaussian distribution with mean ***v*** (called drift rate) and variance ***s*** on every trial. Here, **v** reflects the speed of evidence accumulation and ***s*** the system’s noise, which scales all other parameters and is usually fixed (here to 0.1). The step size for all models was set to 1ms. Responses are triggered when the diffusion reaches a criterion (boundary, determined by the free parameter ± ***a***). We assumed that there was no bias in response selection over the task, because we had an equal number of left and right responses. However, individual trials were allowed to start with a random bias towards one response (start-point variability, parameter ***sz***). We used symmetrical boundaries which were defined as left-hand responses when the positive boundary was reached first, and as right-hand responses when the negative boundary was reached first (see Figure 1B for more details). The non-decision time was modelled as another free parameter ***T_er_***. The stages of the model per trial were defined as a zero-mean baseline until flanker onset, a noisy diffusion with v = 0 during the non-decision time, a diffusion driven by the flanker direction between flanker and target onset (83 ms) with drift = vt × ft, and the target phase thereafter with drift = ***vt*** and the direction of the target. Single-trial values ***vt*** and ***ft*** were determined according to Gaussian variance parameters ***sv*** and ***sf***. For display purposes, in Figure. 2F, we modelled a consecutive return of the decision variable to baseline similar to an Ornstein-Uhlenbeck process to facilitate comparisons between model and BPL. To speed up the model fitting procedure, we did neither simulate baseline periods nor return to baseline during the fitting because these have no effect on model predictions. We compared four different variants of the DDM by fitting their parameters to RT and accuracy data observed in the group of 863 participants using quantile maximum likelihood statistics (Heathcote et al., 2004) and differential evolution algorithms (Price et al., 2005). Additionally, we used a mixture model assuming 2% outliers that were distributed uniformly over the full range of RTs in correct and error responses. This downweighs the impact of possible outliers on model parameters. DDM model 1 used the same drift rate during flanker and target processing, thus not allowing for suppression of distractors (parameter ***f*** fixed to 1). DDM model 2 fixed parameter ***f*** which suppressed flanker processing when below the value of 1. DDM model 3 furthermore included trial-by-trial variance in ***f*** modelled as a zero-mean Gaussian distribution with variance ***sf***. The fourth DDM model extended DDM model 3 to allow for a dynamic decision boundary collapse according to a Weibull distribution scaled by the free parameter ***k***. Here, the dynamic boundary ***u*** at time ***t*** was calculated as follows:

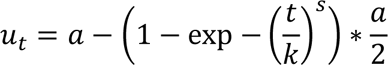

In this equation, ***α*** represents the initial boundary value and ***k*** scales the Weibull distribution (see Supplementary Figure 1 for illustration of the impact of this parameter on boundary dynamics). The shape parameter ***s*** was fixed at 3, to impose a “late collapse” decision strategy. This value/shape was informed by results from a previous study in a large human sample (Hawkins, Forstmann, et al., 2015).

For model comparison, we first computed approximated BIC values (aBIC; akin to White et al. (2011)). Next, we used these individual aBIC values to computed protected exceedance probabilities, which gives the probability that one model was more likely than any other model of the model space (Rigoux et al., 2014).

To simulate the temporal evolution of the modelled diffusion signal depicted in Figure 3 we used the mean maximum likelihood parameters from the group fit obtained for DDM 4. For this simulation, we computed 5000 simulated trials.

### EEG processing

EEG was recorded at 500Hz from 60 Ag/AgCl sintered electrodes with A1 as reference channel and an electrode placed over the sternum as the ground electrode arranged in the extended 10-20 system using BrainAmp amplifiers (Brain Products, Gilching, Germany). Impedances were kept below 5 kΩ. Pre-processing of the EEG data was done under Matlab 2021b (The MathWorks, Natick, MA) and the EEGlab 13 toolbox (Delorme & Makeig, 2004) using custom routines as described previously (Fischer et al., 2018). Pre-processing steps included: (1) Filtering (0.5 Hz high- and 42 Hz low-pass filter), (2) re-referencing to common average, (4) Segmentation into stimulus-locked epochs spanning −1.5 to 2 s relative to target onset, (5) automatic epoch rejection. Here, epochs contaminated with artefacts were rejected using a dynamically adjusted rejection threshold to remove at least 1 trial separately for error and correct responses and maximally 5% per condition (average number of rejected epochs: 41, range 11–53). (6) removal of blink, eye-movement, and other, less homogenous artefact components (average number of removed components = 3.4, range 1–14) using adaptive mixture independent component analysis (AMICA; Palmer et al., 2012). Finally, we extracted response-locked epochs spanning from −1 to 1 s from the stimulus-locked epochs.

### EEG analyses

First, we convolved the artefact free response-locked EEG signal, with a series of complex Morlet wavelets between 13 and 25 Hz. We used 20 linear steps and a wavelet width of six cycles. Data were then log-transformed and normalized. When comparing the averaged signal over contralateral (electrodes C3/CP3 for right and C4/CP4 for left) and ipsilateral (electrodes C3/CP3 for left and C4/CP4 for right) motor cortices, we show an overall decrease in beta power around 300ms prior to the response, which was followed by a consecutive increase in beta power (i.e., beta rebound) around 400ms post response (Figure 3c). To confirm the lateralisation of the signal to the hemisphere initiating a response, we used single-trial multiple robust regression. Here, we regressed the factor response hand (−1 = left, 1 = right) onto the convolved signal across the collapsed frequency range of 13–25 Hz for every electrode and time point, while controlling for unspecific task effects. Noise-Regressors included: congruency, flanker distance, RSI, following RSI, trial number, and log transformed RT. This analysis confirmed response related beta band lateralisation, whereby beginning around 100ms before response onset, beta power decreased over the sensorimotor cortex contralateral to the response. This was reflected in a positive covariation between response hand and the signal in the beta band which was maximal at C4/CP4, and a negative covariation at maximal at C3/CP3 (Figure 2a). We chose electrodes with a maximal effect (C3, C4, CP3, CP4) for all further analyses of lateralised beta power (BPL).

To derive BPL we subtracted beta band power over the inactive sensorimotor cortex (i.e., the electrodes side ipsilateral to the hand that gave the response in the trial) from the beta power recorded over the active (contralateral) sensorimotor cortex. This difference signal thus compares the degree of beta power reduction between both hemispheres, presumably reflecting differential motor activation (see Figure 3). To investigate single-trial associations of beta thresholds and RT, we used the response-locked single-trial signal (mean ± 12 ms) and regressed the log-transformed RT onto the beta threshold while accounting for unspecific task effects according to the following equation:

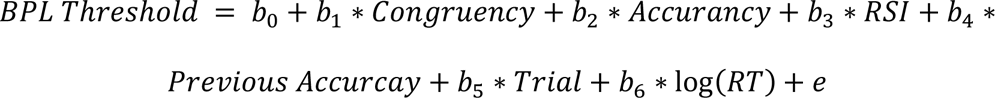

In an explorative analysis, we used the same model to test the separate contributions of each hemisphere (ipsilateral and contralateral beta power).

## Data and code availability

The conditions of our informed consent form do not permit public archiving of the raw data because participants did not provide sufficient consent. Researchers who wish to access processed and anonymized data from a reasonable perspective should contact the corresponding author. Data will be released to researchers if it is possible under the terms of the GDPR (General Data Protection Regulation). The code of the toolbox we used for the regression analysis can be found here: http://www.adrianfischer.de/teaching.html. The code for the DDM models is available on request. Data and analyses scripts of the confirmation sample can be downloaded on the Open Science Framework at [insert after acceptance for publication].

## Acknowledgments

This research was supported by the Deutsche Forschungsgemeinschaft (KL 2337/2-1 to TAK, UL 196/3-3 to MU) and by the European Research Council (ERC) under the European Union’s Horizon 2020 research and innovation programme (ERC advanced grant agreement No 101018805 to MU).

## Supplements

### Additional info on DDMs

**Supplementary Table 1-.** Maximum likelihood parameters of three best fitting DDMs. The table shows the parameters obtained by fitting the models to the individual subject data. DDM3 was comprised of the standard-parameters reflecting drift rate (v), flanker weighting (f), boundary (a), and non-decision time (Ter), and variance parameters that determined how much standard-parameters varied from trial-to-trial (sv, sz, st, sf). DDM4 extended DDM3 to allow for a dynamic decision boundary collapse according to a Weibull distribution scaled by the free parameter k. In DDM5 the free parameter k is used as a multiplicative gain (urgency signal) instead of boundary collapse. In this model bounds are fixed but evidence accumulation is subject to a gain signal that increases with the duration of the decision. See Supplementary Figure 2 for examples of dynamic bound models and their relation to the scaling parameter k. In all models, diffusion noise (s) was fixed to 0.1 as commonly seen in the literature (e.g., White et al., 2011). To translate Ter values and their variance into ms they must be multiplicated by 1000 and 100 respectively. We set the bias parameter for start points (z) are set to zero for all models because the task included exactly 50% left- and right-hand responses.

| Parameter | DDM3<br>Mean (SD) | DDM4<br>Mean (SD) | DDM5<br>Mean (SD) | boundary parameters<br>(i.e., boundary of flat prior) |
| --- | --- | --- | --- | --- |
| Drift rate (**v**) | 4.54 (1.42) | 4.23 (1.46) | 4.17 (1.48) | 0.01 - 8.5 |
| Variance in drift rates (**sv**) | 0.79 (0.54) | 0.80 (0.54) | 0.83 (0.53) | 0 - 1.5 |
| Boundary (**a**) | 0.37 (0.08) | 0.37 (0.07) | 0.37 (0.08) | 0.01 - 0.45 |
| Variance in start points (**sz**) | 0.19 (0.09) | 0.17 (0.09) | 0.18 (0.09) | 0.05 - 0.3 |
| Non-decision time (**T<sub>er</sub>**) | 0.29 (0.02) | 0.29 (0.02) | 0.29 (0.02) | 0.1 - 0.4 |
| Variance in T<sub>er</sub> (**st**) | 1.14 (0.41) | 1.15 (0.41) | 1.15 (0.43) | 0 - 2 |
| Flanker suppression (**f**) | 0.48 (0.25) | 0.47 (0.27) | 0.49 (0.28) | 0.1 - 1.5 |
| Variance in **f** | 0.70 (0.29) | 0.75 (0.27) | 0.75 (0.28) | 0 - 1.5 |
| Boundary collapse / gain parameter (**k**) | - | 2.41 (0.60) | 2.49 (0.59) | 10e-4 - 3 |
| (fixed) noise of within-trial diffusion (**s**) | 0.1 | 0.1 | 0.1 | 0.1 |
| (fixed) average start point bias (**z**) | 0 | 0 | 0 | 0 |

**Supplementary Figure 1.**
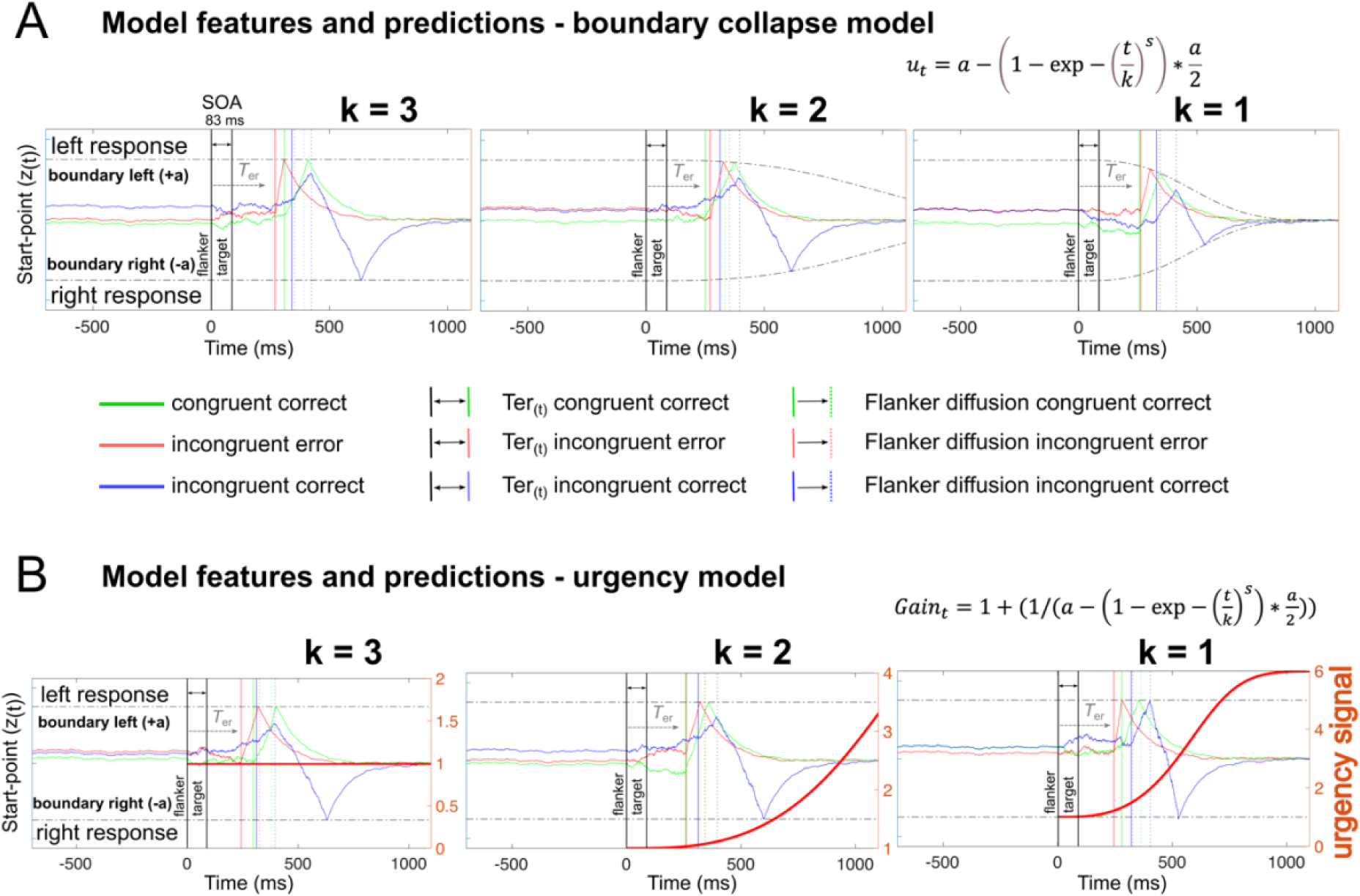
Illustration of different within-trial bound dynamics. **(A)** Three illustrative boundary collapse trajectories depending on the scaling parameter k. Here, smaller values are associated with faster boundary collapse. **(B)** Depicts 3 different urgency signal trajectories. The urgency signal (orange line) is used as a multiplicative gain of the diffusion and noise. As for the boundary collapse models, lower values of the free parameter k are associated with a stronger urgency signal. All trial simulations used the fitted group means (see Supplementary table 1) except the dynamic decision bound scaling factor that was set to either 1, 2 or 3. For model fitting we applied the following hard priors for k, which can be seen as boundary parameters: [10e-4 – 3].

### Urgency model details

We also considered a different kind of dynamic decision boundary model. In this model, collapsing decision boundaries are facilitated via an urgency signal (Cisek et al., 2009; Hanks et al., 2014). Here, an urgency signal that increases with decision time is used as a multiplicative gain of the diffusion and noise (see Supplementary Figure 1 and 2A for an illustration). The urgency signal (Gain) at a given time within a trial (t) was given by:

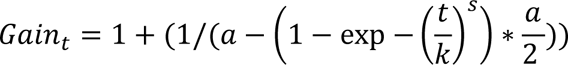

where α represents the initial boundary value and *k* scales the urgency signal according to an inverted Weibull distribution with the shape parameter ***s*** fixed at 3.

Model comparison (Supplementary Figure 2B) and validation (Supplementary Figure 2C) revealed that our data is similarly well captured by the urgency (DDM5) and collapsing bound model (DDM4). However, when considering the trough of movement-selective beta suppression around response time as proxy for decision boundary thresholds, both dynamic decision boundary models make distinct predictions. While the collapsing bound model predicts that slower responses are made at lower decision boundary thresholds, the urgency model would predict similar decision boundaries for slow vs. fast responses (Supplementary Figure 2D). Hence, based on our robust finding, that the trough of the movement-selective beta suppression around response time is scaling with RT, we consider the collapsing bound model as more neural plausible.

**Supplementary Figure 2.**
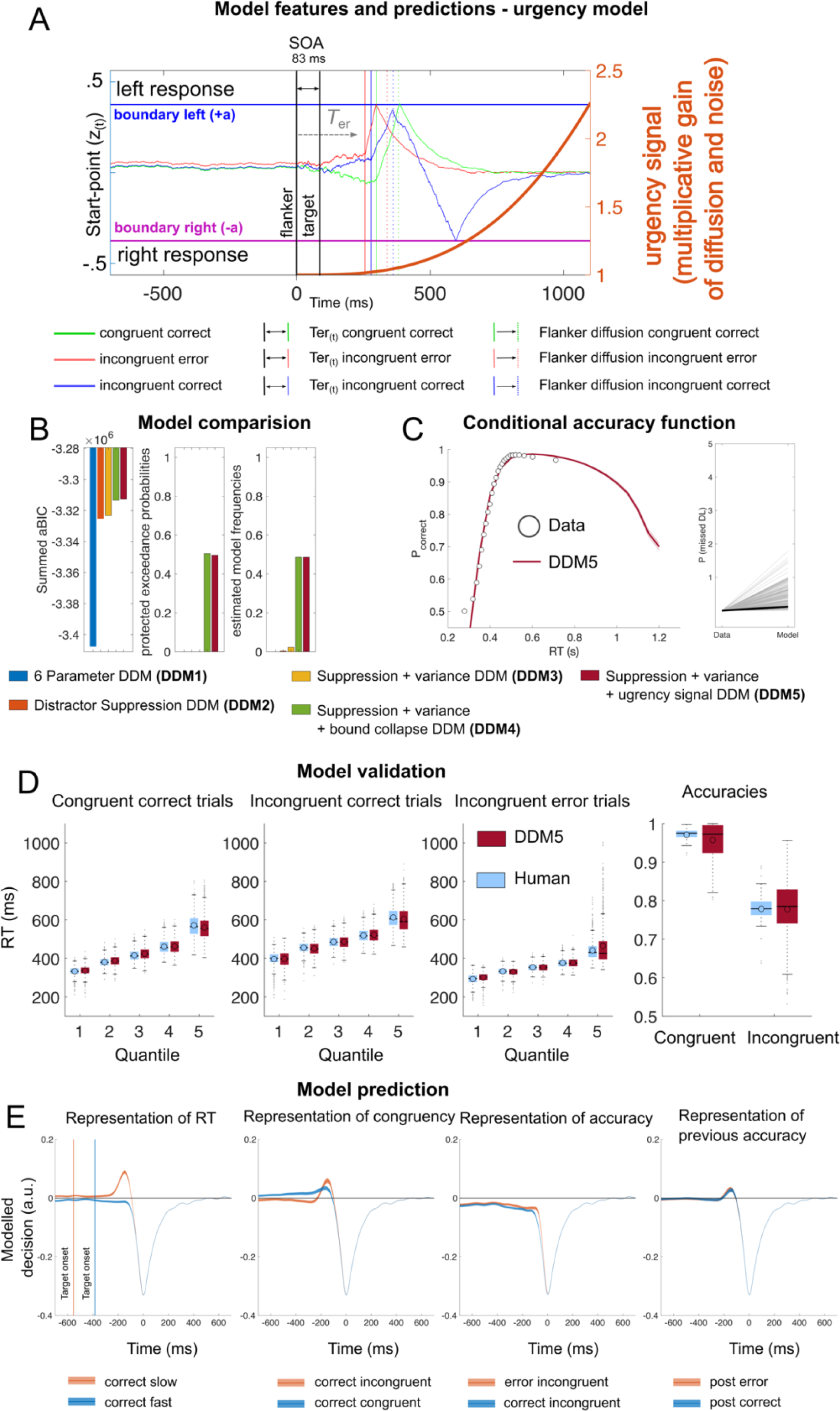
Model features, fit, and predictions of the urgency model. **(A)** Illustration of the multi-stage DDMs based on simulated single trial decision processes. Features of this model are the same as described in the main manuscript (Figure1 B), with the exception that here the free parameter k is used to scale the urgency signal (orange line) that increases with decision time. This urgency signal is used as a multiplicative gain of the noise and diffusion. **(B)** Cumulated approximated Bayesian Information Criterion (aBIC) scores over participants of each candidate model. Higher values indicate a better fit of the models to the behavioral data. Results indicate that the dynamic bound model (DDM4) and the urgency model (DDM5) provides a similar fit to the data. Protected exceedance probabilities (pEP) draw a similar picture. **(C)** depicts Conditional accuracy functions and proportion of missed deadlines. Points indicate mean accuracy of trials sorted by RT into 25 equal-sized bins informed form the empirical data. Line shows best fits of piece-wise logistic regressions to each subject’s modelled single-trial data. Shades indicate ± s.e.m. Note that for model fitting we excluded all missed trials from the empirical data, hence the ground truth for missed deadlines of the data is zero. **(D)** shows RT quantile fits and accuracy fits of the urgency model against human data. This model validation is suggesting a good fit to the data. **(D)** To characterise predictions of the urgency DDM we simulated 5000 individual trials using the mean of the best fitting individual parameters (see Supplementary Table 1). **(E)** shows the modelled decision process locked to response separated by response speed, congruence, accuracy, and previous accuracy. These simulations predict that decision thresholds are unaffected by these task factors.

**Supplementary Figure 3.**
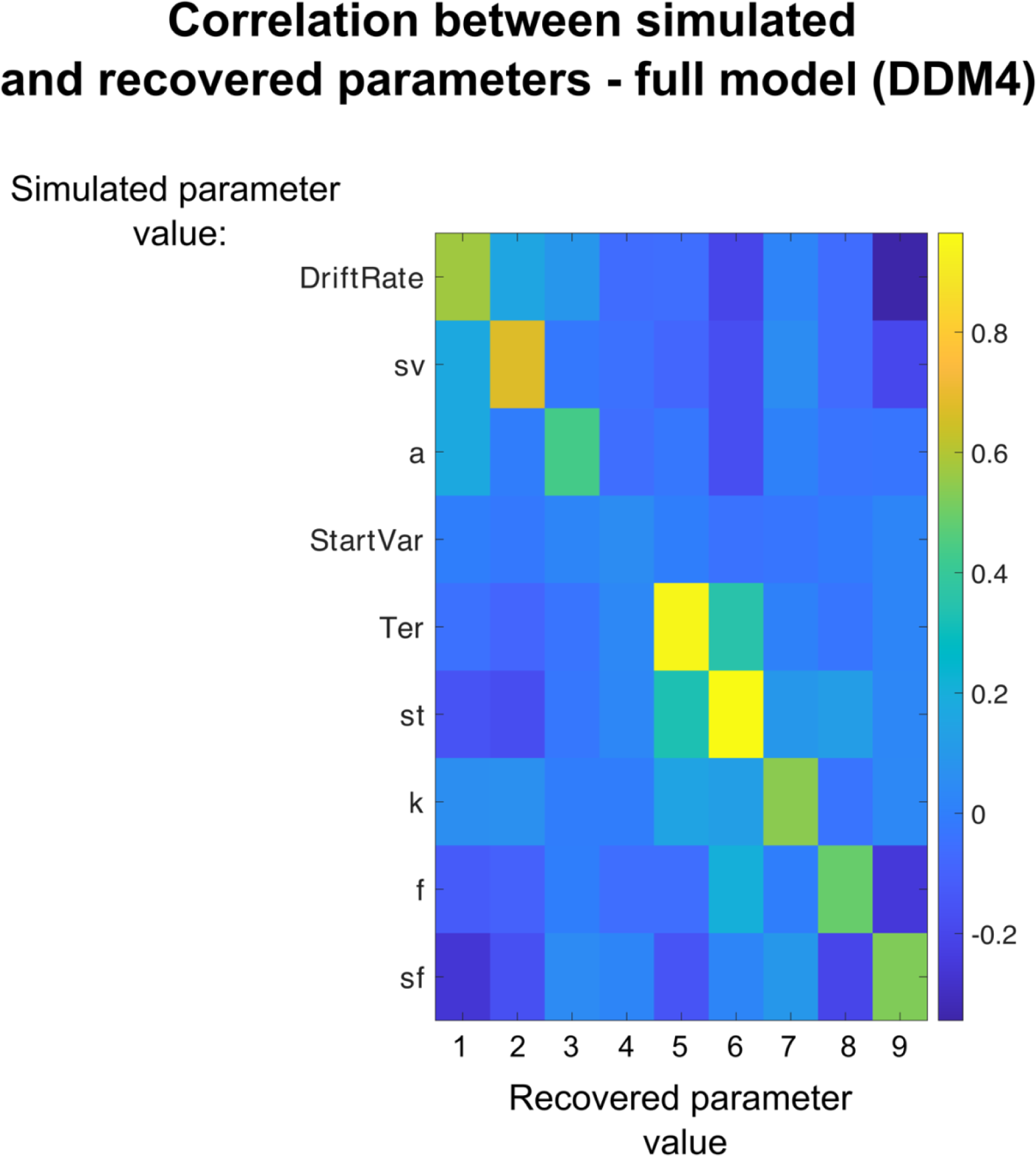
Parameter recovery DDM4. For parameter recovery analyses, we randomly drew model parameters out of a Gaussian distribution with mean and variance equal to the observed fitted parameters across the whole group to reduce parameter value combinations that were extremely unlikely to occur in human data. We simulated 1.000 parameter combinations and used the same differential evolution algorithm to recover the fitted models. Models that produced no errors at all or for which constraints were not met, were discarded from analysis. As in the human data, we used 5.000 trials per simulation. Plotted are correlations between simulated and recovered parameters for the full model (DDM4). Results indicate that parameter values that were used to simulate data from the full feature DDM (ordinate) tended to correlate with the parameter values best fit to those synthetic datasets (abscissa). *Note:* the full feature DDM consisted of nine parameters: drift rate (v), variance in drift rates (sv), boundary (a), variance in start points (sz), non-decision time (Ter, reflecting visual processing and motor execution times), variance in Ter (st), bounds collapse (k), flanker suppression (f), and variance in f (sf).

### Supplementary EEG results

**Supplementary Figure 4.**
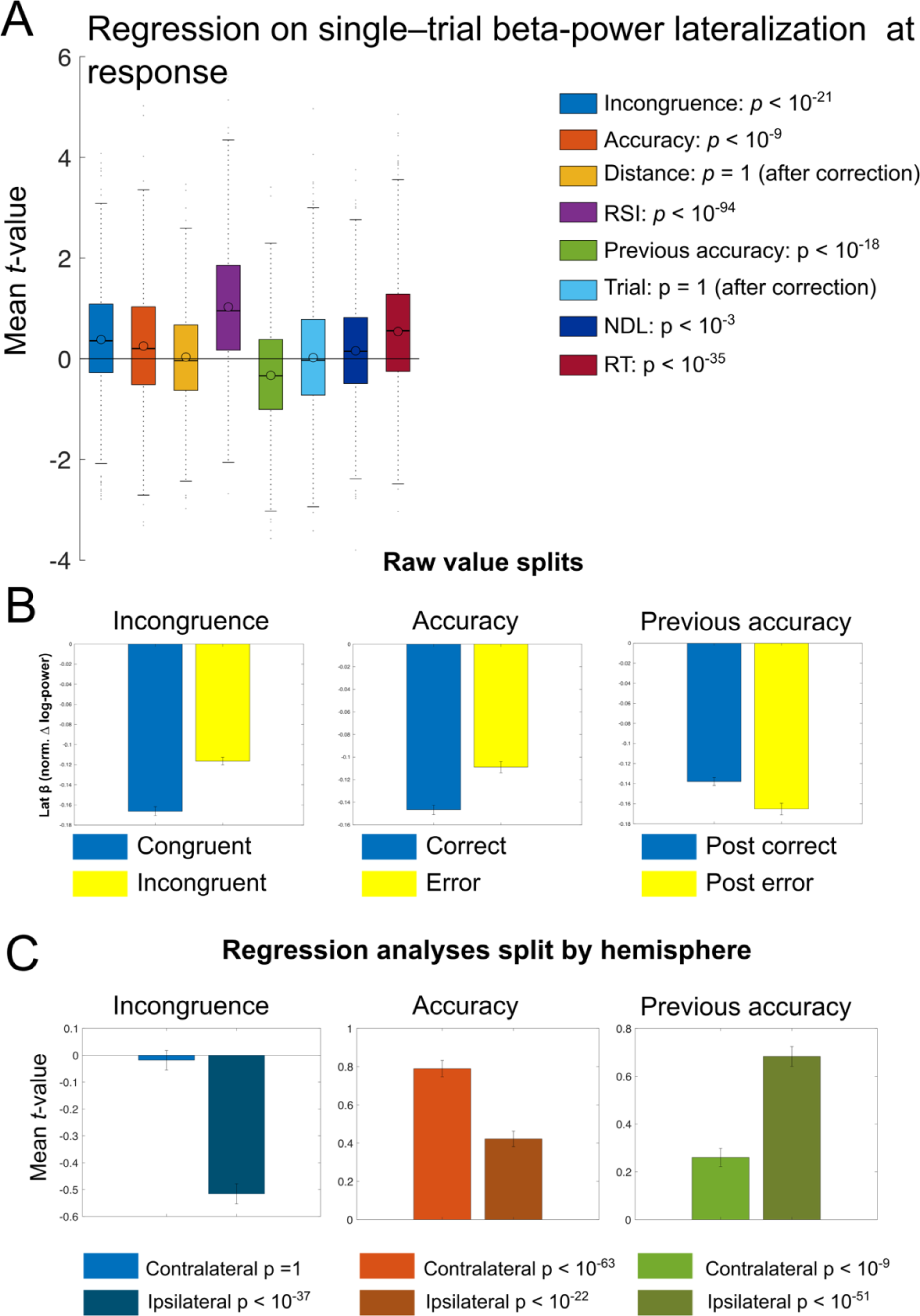
Single-trial regression on beta lateralisation at response. **(A)** Results of a single-trial regression analysis on BPL at response (mean ± 12ms surrounding button press), comparing relevant task factors. **(B)** Shows raw values splits for Incongruence (congruent vs incongruent), Accuracy (correct vs error) and Previous Accuracy (post correct vs post error). **(C)** Display results of separate regression analyses split by ipsilateral (hemisphere that does not induce the choice on the current trial) and contralateral hemispheres. Results indicate, that incongruent trials cause reduced BPL (t(862) = 10.21, p < .001, d = 0.35, CI [0.31, 0.45], BF_10_ > 100). This effect is driven by changes over the ipsilateral hemisphere where beta power was more strongly reduced on incongruent trials. This may reflect the pre-activation of the competing response tendency. Similarly, errors were associated with reduced BPL (t(862) = 6.52, p < .001, d = 0.22, CI [0.17, 0.32], BF_10_ > 100). This result is partly driven by stronger increases in beta power of the contralateral hemisphere. However, it should be noted that this effect is not evident, when comparing RT matched BPL on correct vs error trials (see Figure 3F), which makes this effect difficult to interpret. Following errors, BPL is larger (more negative, t(862) = −9.34, p < .001, d = −0.32, CI [−0.4, −0.26], BF_10_ > 100) which is caused by a stronger increase in BP over the ipsilateral hemisphere. This may reflect static decision policy adaptations to increase accuracy after errors (Fischer et al., 2018; Murphy et al., 2016). An additional interpretation is that following errors, the unchosen response is more effectively suppressed. Error-bars reflect 99.9% CI, **(A and C)** depicts mean within participants t-values, and statistics are results of t-tests of individual within-subject regressions against zero. NDL reflects the distance from the last break in the task, and trial the general time on task, together RSI these regressor were used as noise regressors.

**Supplementary Figure 5.**
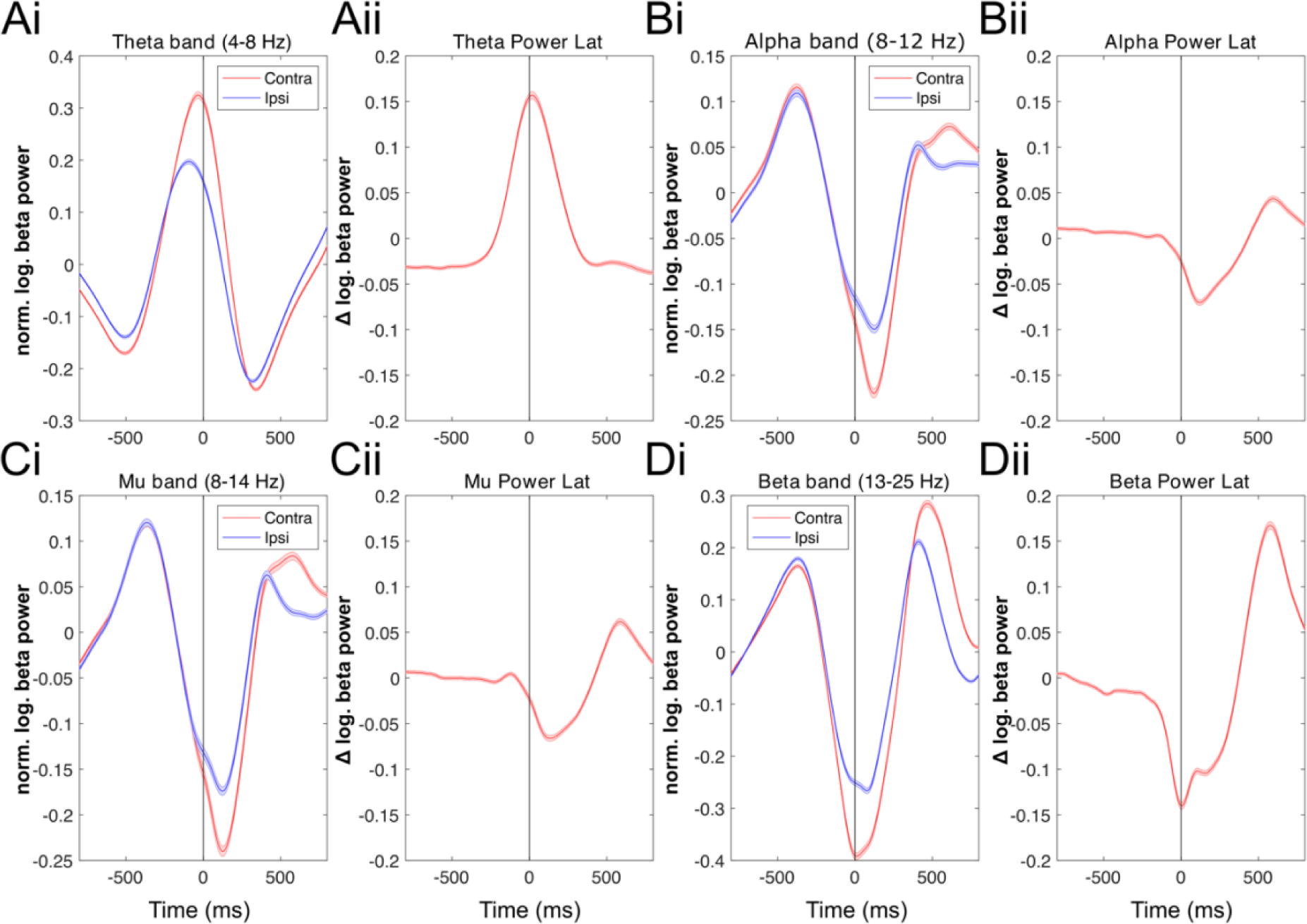
Movement selective lateralization signals across frequency bands. **(A-D)** demonstrate that lateralization signals across frequencies is composed of the activity-difference recorded over contralateral and ipsilateral motor cortices. We chose electrodes with a maximal effect (C3, C4, CP3, CP4) for analyses of lateralised signals. Specifically, we subtracted the power in the respective band over the inactive sensorimotor cortex (i.e., the electrodes side ipsilateral to the hand that gave the response in the trial) from the beta power recorded over the active (contralateral) sensorimotor cortex. This difference signal thus compares the degree of power reduction between both hemispheres, presumably reflecting differential motor activation. Results indicate that the effect is most pronounced in the beta band frequency.

**Supplementary Figure 6.**
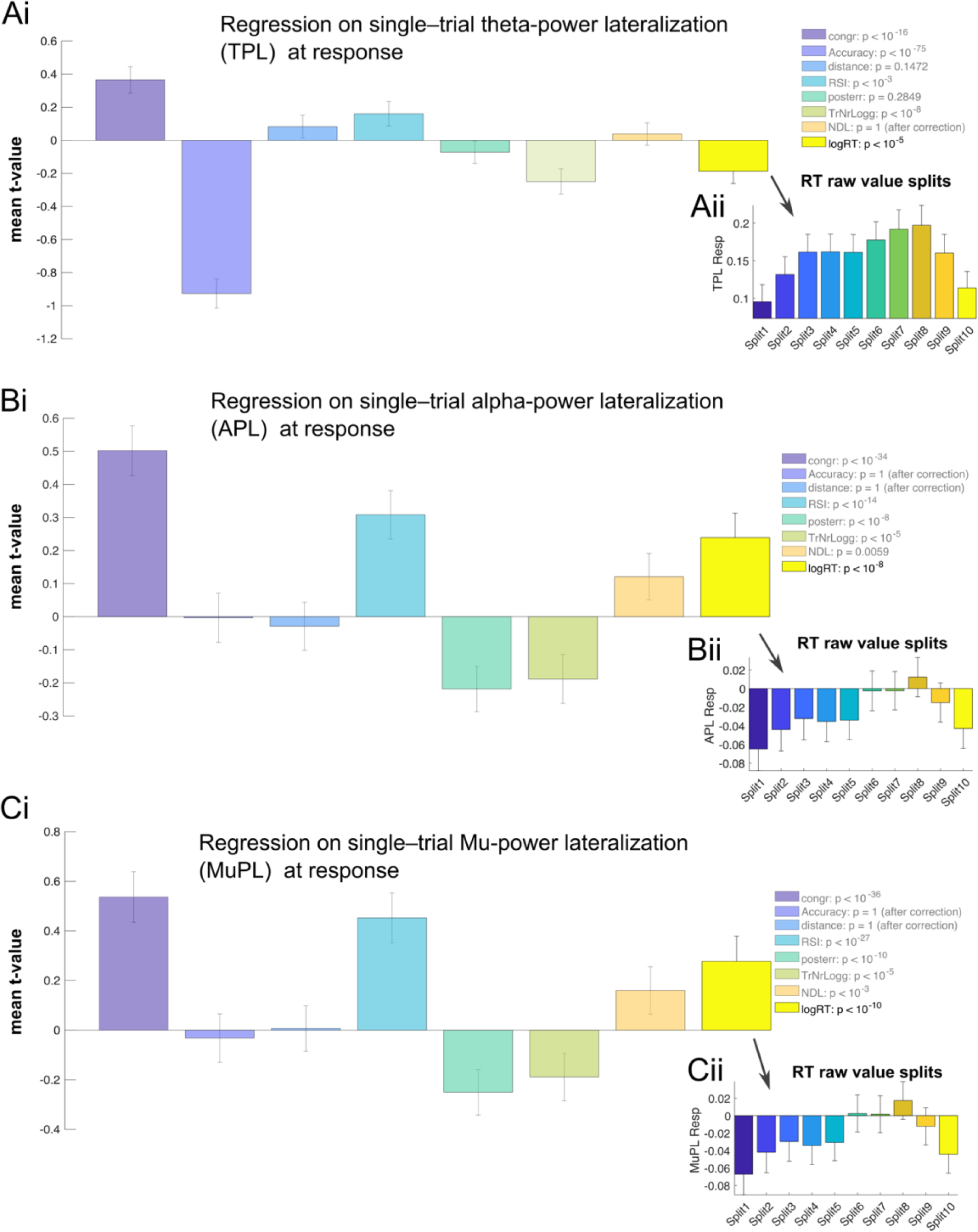
Single-trial regression on different movement selective frequency band lateralisation signals at response. To investigate single-trial associations of movement selective frequency band suppression around response and RT, we used the respective lateralised response-locked single-trial signal (mean ± 12 ms) and regressed the log-transformed RT onto the threshold while accounting for unspecific task effects. Results indicate that that both APL (8-12 Hz) and MuPL (8-14 Hz) is decreasing (i.e., boundary collapse) with increasing RTs (thus representing a decision threshold decrease with increasing decision time). Moreover, an opposite effect was present for lateralization in the theta frequency band (4–8 Hz). Error-bars reflect 99.9% CI.

### Confirmation sample

To replicate the key findings in an independent sample, we analysed data of one hundred nineteen subjects (mean age 22.89 years (range: 18 - 38), 70 females). These datasets were collected at the Otto-von-Guericke University Magdeburg as part of an ongoing genetic association study using the same approved study protocol as described in the main manuscript. Behavioural analyses of this sample confirmed that general behaviour effects were in accordance with typical results seen in flanker tasks. They reflected interference effects, with slower (ΔRT = + 42 ms, t(118) = −43.25, p < .001, d = 1.6986, BF_10_ > 100) and more erroneous (Δaccuracy = 19%, t(118) = 34.51, p < .001, d = 3.7163, BF_10_ > 100) responses on incongruent trials. The overall accuracy was 86% (SD=3.94). To replicate key findings, we used the same analyses procedures as for the main sample. Results replicate the findings for the main sample: First, we show that the dynamic decision boundary model outperformed static decision boundary conflict DDMs and captured conflict related RT and accuracy well (see Supplementary Figure 7). Moreover, the EEG analysis validated our previous conclusion that the trough of movement-selective beta suppression around response time may reflects a collapsing decision bound (see Supplementary Figure 8).

**Supplementary Figure 7.**
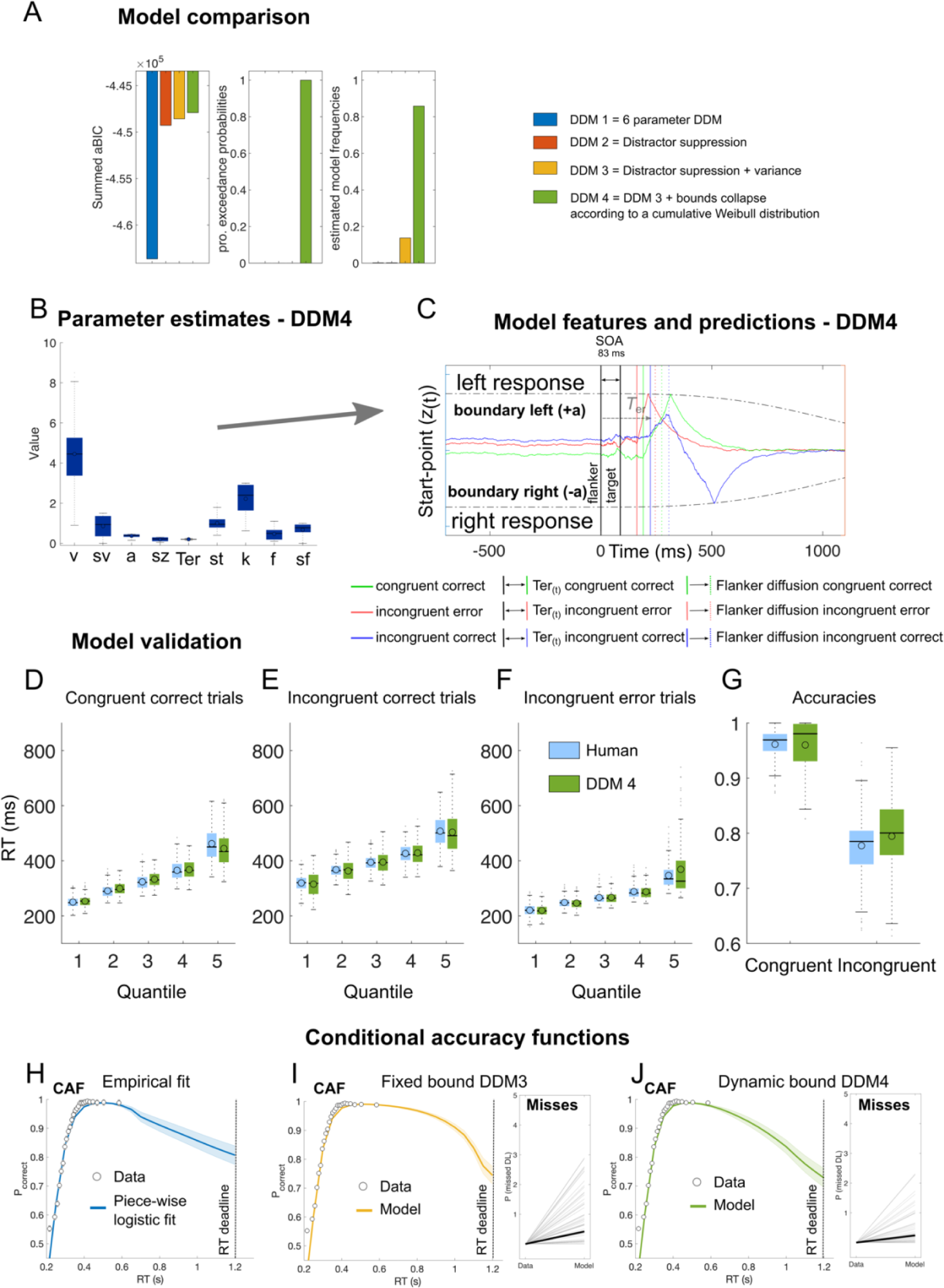
Model comparison and validation in the confirmation sample. **(A)** Cumulated approximated Bayesian Information Criterion (aBIC) scores, protected exceedance probabilities (pEP) and estimated model frequencies indicate that the dynamic bound model (DDM4) provides the best fit to the data. **(B)** The drift-diffusion model (DDM) included nine free parameters. In addition to the standard-parameters reflecting drift rate (v), flanker weighting (f), boundary (a), and non-decision time (Ter), and variance parameters that determined how much standard-parameters varied from trial-to-trial (sv, sz, st, sf), the dynamic boundary model included the free parameter k that scaled the boundary collapse that increases with the duration of the decision. **(C)** shows single trial simulations and model features based on the mean of the group best fit parameters displayed in **(B). (D–G)** shows quantile fits of the model against human RT data and **(F)** shows model and human accuracy. In all conditions (congruent & incongruent correct as well as incongruent error), the model captures the RT data in each quantile, suggesting a good fit to the data. *Note*: boxes = interquartile range (IQR), o = median, - = mean, whiskers =1.5 × IQR, grey dots = outlier. **(H)** Shows the conditional accuracy function. Here, points indicate mean accuracy of trials sorted by RT into 25 equal-sized bins. Line shows best fits of piece-wise logistic regressions to each subject’s single-trial data. Shades indicate ± s.e.m. **(I-F)** depicts Conditional accuracy functions and proportion of missed deadlines from the full fixed bound model (DDM3; **K**) and the collapsing boundary model (DDM4; **L**). Points indicate mean accuracy of trials sorted by RT into 25 equal-sized bins informed form the empirical data. Line shows best fits of piece-wise logistic regressions to each subject’s modelled single-trial data. Shades indicate ± s.e.m. Note that for model fitting we excluded all missed trials from the empirical data, hence the ground truth for missed deadlines of the data is zero.

**Supplementary Figure 8.**
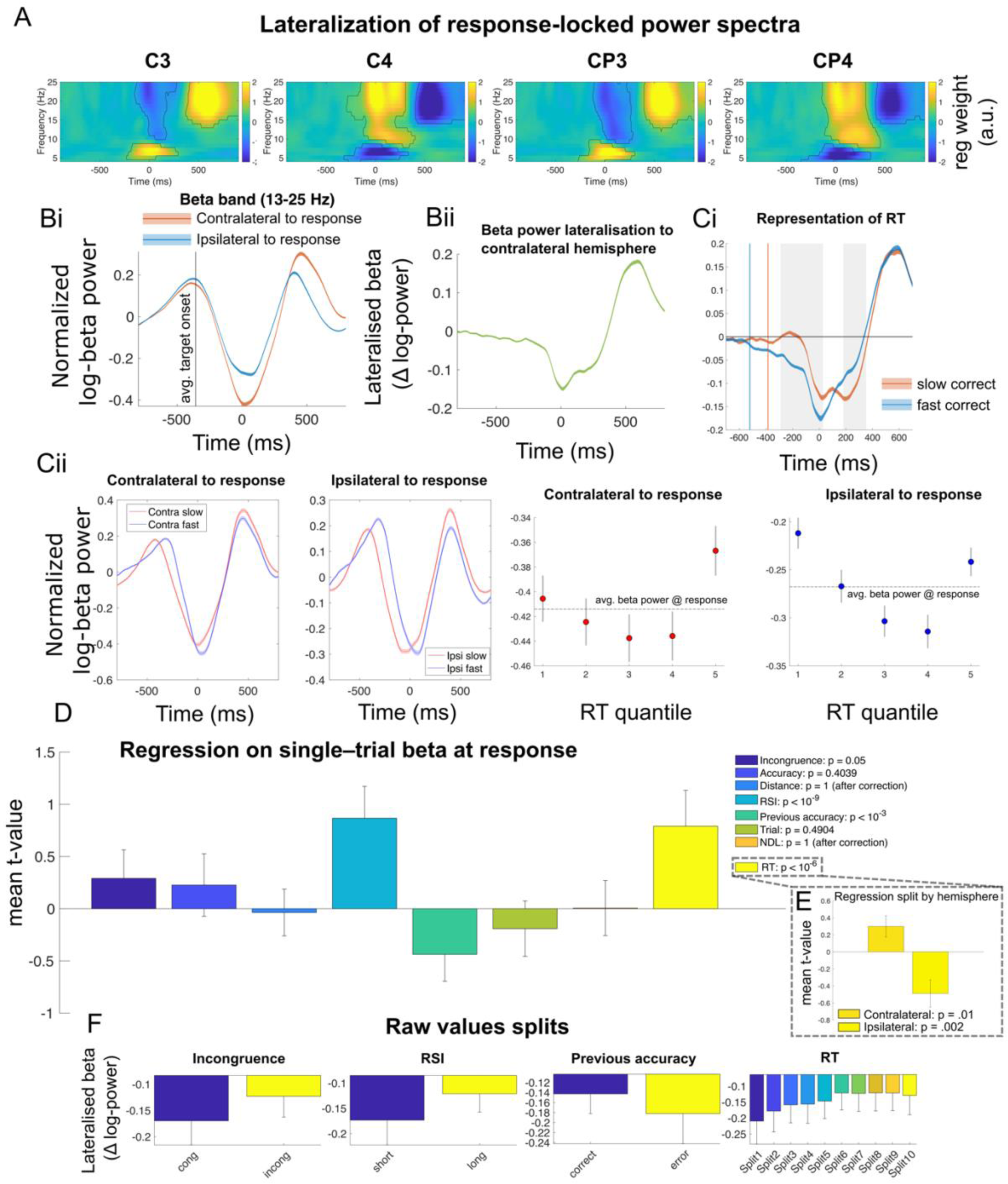
EEG results confirmation sample. **(A)** Shows the result of a single-trial regression analysis comparing responses with the left and right hand, while controlling for congruency, flanker distance, response stimuli intervals (RSI), following RSI, trial number, and log transformed RT. Data suggests, that Beta power more strongly decreases when a response is given with the contralateral compared to the ipsilateral hand. This effect was most pronounced over the electrodes C3/4 and CP3/4. **(B)** Beta power lateralization signal **(Bii)** is composed of the activity-difference recorded over contralateral and ipsilateral motor cortices **(Bi)**. **(Ci)** Beta power lateralization around response time may reflects collapsing decision thresholds, whereby faster responses (based on a median split) are made at higher thresholds. This effect is partly explained by a reduction of beta power over the contralateral hemisphere **(Cii, first left)**, while on average beta power over the ipsilateral hemisphere is barely changed when responses are given later in a trial **(Cii, second to the left).** Yet, BPL showed a stronger influence of response time than either of the hemispheres alone suggesting that it may be an especially valid marker of response threshold modulations. Boxes next to the average beta power traces show beta power (mean ± 12ms surrounding button press) divided into five equally sized (within subject) RT bins. * = uncorrected significant difference to mean beta power (dashed lines) **(D)** Results of a single-trial regression analysis on BPL at response (mean ± 12ms surrounding button press). This analysis confirms collapsing decision boundaries on a single-trial level while controlling for other task factors that potentially influence BPL. **(E)** Display results of separate regression analyses on beta power split by ipsilateral (hemisphere that does not induce the choice on the current trial) and contralateral hemispheres. Results indicate that RT dependency in both hemispheres (contralateral: mean t-value (SD): 0.29 (0.12), t(118) = 2.39, p = 0.01, d = 0.22, CI [0.05, 0.54]; ipsilateral: −0.48 (0.16), t(118) = −3.05, p = 0.002, d = −0.28, CI [−0.80, −0.17]). However, the effect is stronger when the relative signal per trial is investigated, than when either of the hemispheres beta power is analysed alone (BPL effect: 0.61 (0.13), t(118) = 4.85, p = 3.7931e-06, d = 0.45, CI [0.36, 0.86]). These data suggest that peak BPL, coinciding with the motor response, may be an especially valid marker of response threshold modulations. **(F)** BPL raw values at response time plotted against GLM factors. In addition to decreases in BPL with increasing RT, we found a significant difference in BPL at response execution between congruent and incongruent trials with higher BPL for congruent trials, which might be interpreted as enhanced motoric readiness caused by the distractor stimuli. Moreover, we found that BPL at response execution was increased on post-error trials, which may reflect post-error adaptions (Fischer et al., 2018). Finally, longer response stimuli intervals (RSIs) were associated with decreased BPL. *Note.* Contours in **(A)** represent significant cluster after cluster based permutation testing (Maris & Oostenveld, 2007). Error-bars and shades reflect 99.9% CI, **(D)** depicts mean within participants t-values, and statistics are results of t-tests of individual within-subject regressions against zero. NLD reflects the distance from the last break in the task, and trial the general time on task, together RSI these regressor were used as noise regressors.

